# Ablation of *OCT4* function in cattle embryos by double electroporation of CRISPR-Cas for DNA and RNA targeting (CRISPR-DART)

**DOI:** 10.1101/2023.07.07.548144

**Authors:** Jada L. Nix, Gustavo P. Schettini, Savannah L. Speckhart, Alan D. Ealy, Fernando H. Biase

**Affiliations:** School of Animal Sciences, Virginia Polytechnic Institute and State University, Blacksburg, VA

**Keywords:** CRISPR, electroporation, gene editing, preimplantation development, POU5F1

## Abstract

CRISPR-Cas ribonucleoproteins are important tools for gene editing in pre-implantation embryos. However, the inefficient production of biallelic deletions in cattle zygotes has hindered mechanistic studies of gene function. In addition, the presence of maternal RNAs that support embryo development until embryonic genome activation may cause confounding phenotypes. Here, we aimed to improve the efficiency of biallelic deletions and deplete specific maternal RNAs in cattle zygotes using CRISPR-Cas editing technology. Two electroporation sessions with Cas9D10A ribonucleoproteins targeting exon 1 and the promoter of *OCT4* produced biallelic deletions in 91% of the embryos tested. In most cases, the deletions were longer than 1000 nucleotides long. Electroporation of Cas13a ribonucleoproteins prevents the production of the corresponding proteins. We electroporated Cas9D10A ribonucleoproteins targeting exon 1, including the promoter region, of *OCT4* in two sessions with inclusion of Cas13a ribonucleoproteins targeting *OCT4* mRNAs in the second session to ablate *OCT4* function in cattle embryos. A lack of *OCT4* resulted in embryos arresting development prior to blastocyst formation at a greater proportion (13%) than controls (31.6%, P<0.001). The few embryos that developed past the morula stage did not form a normal inner cell mass. Transcriptome analysis of single blastocysts, confirmed to lack exon 1 and promoter region of *OCT4*, revealed a significant (FDR<0.1) reduction in transcript abundance of many genes functionally connected to stemness, including markers of pluripotency (*CADHD1*, *DPPA4*, *GNL3*, *RRM2*). The results confirm that *OCT4* is key regulator of genes that modulate pluripotency and is required to form a functional blastocyst in cattle.

**Significance Statement:** CRISPR-Cas mediated DNA editing can revolutionize agriculture and biomedicine due to its simplicity of design and use. Modifications induced in embryos, though challenging to accomplish, are beneficial for the advancement of livestock production and the study of biological function. Here, we developed an approach using CRISPR-Cas enzymes to remove DNA segments of the cattle genome in one-cell embryos. Our results show major advancement in the efficiency of producing large deletions in the genome of cattle embryos. Using our approach, we removed the function of the *OCT4* gene. Our results confirmed *OCT4* as a major regulator of pluripotency genes during embryo development and its requirement for the formation of an inner cell mass in cattle.

## Introduction

The driving force behind gene functionality studies is the targeted alteration of genomic sequences followed by observation of phenotypic deviations. The deletion of functional sequences in the genome, also called knockouts (KO), can be used to study the roles of genes during pre-implantation embryonic development (1). Mechanistic studies of gene function provide information connecting genome and phenotype during early embryogenesis, and the data may be used to better understand biological function (2–5) or disease (6–8). The CRISPR-Cas system has been the method of choice for most researchers wishing to alter genome sequences in somatic (9–11), germ (12–15), or embryonic cells (16–25). CRISPR-Cas systems have gained traction due to the simplicity of design and synthesis of gRNAs with sequence complementarity to the target region (26) and improved efficiency when compared to other common methods for sequence alterations (27–30).

Despite recent advancements in protein engineering giving rise to CRISPR-Cas ribonucleoproteins of greater efficiency and specificity (31), biallelic deletion efficiency, or the deletion of targeted sequences in both chromosomes, remains low in CRISPR-Cas treated zygotes across many species, including cattle (32–37). Interestingly, only four reports provide data on biallelic deletion efficiency in studies utilizing CRISPR-Cas introduced through electroporation of cattle zygotes (33, 35, 38, 39). These studies averaged 75% of sampled embryos containing partial deletions, with the presence of at least one wildtype allele, and 59% containing full deletions with no wildtype alleles. Some intrinsic factors of zygote biology, such as chromatin compaction and the timing of DNA replication, may impair deletion efficiency due to sequence inaccessibility for CRISPR-Cas binding or the increased number of target sites requiring DNA cleavage. Though the introduction of increased amounts of CRISPR-Cas by more intense electroporation conditions is shown to improve editing efficiencies in cattle zygotes, embryonic mortality increases in tandem (40). Alternate methods for increasing CRISPR-Cas content in the zygote have been used, such as zona pellucida drilling prior to electroporation in cattle (35) or zona removal in swine (41). These methods may improve CRISPR-Cas delivery but do not mitigate the setback of embryo mortality. Additionally, it has been suggested that maternally inherited mRNA, present in mammalian zygotes (42–45), may support sufficient protein production in the absence of a functional gene. The presence of mRNA resulting from the gene of interest likely hinders gene functionality studies in preimplantation embryos and may be responsible for inconsistent knockout phenotypes. To that end, Cas13a (46) may be used to knockdown maternal or nascent mRNA and further obstruct protein production, but this element has not been accounted for in previous cattle studies. Altogether, many factors can influence the efficiency of CRISPR-Cas systems in pre-implantation embryos.

The gene *OCT4*, or octamer transcription factor 4, is thought to maintain pluripotency in early cattle (47, 48) and human (49, 50) embryos through its role as a transcription factor for many pluripotency related genes (49, 51). Additionally, it functions in the HIPPO signaling pathway (52) and is thought to be a key regulator of the first cell lineage differentiation event in cattle. The function of OCT4 has been studied in murine preimplantation embryogenesis models, and these studies show that normal blastocyst development and first cell lineage differentiation are possible in the absence of an *OCT4* gene (53, 54), but one murine model results in the development of blastocysts with absent inner cell mass (49). As HIPPO signaling processes vary between bovine and murine preimplantation development (52, 55), these results may not provide adequate translation of information regarding human cell lineage differentiation. Studies to determine the role of *OCT4* have been completed by CRISPR-Cas mediated KOs in cattle zygotes, but these studies produced varying outcomes and inconsistent phenotypes (35, 36, 56, 57). Most studies report *OCT4* KO cattle embryos maintaining the ability to reach the blastocyst stage and effectively completing the first cell lineage differentiation event in the absence of this gene (38, 58, 59). Conversely, one report showed developmental arrest at the morula stage, prior to cell lineage specification (35). This variability may be due to unaccounted factors, such as maternal or pre-existing mRNA, the common presence of wildtype alleles in CRISPR-Cas genome edited cattle zygotes, and how zygotes were generated.

Here, we aimed to improve the efficiency of CRISPR-Cas mediated biallelic deletions in cattle zygotes while degrading preexisting RNAs transcribed from the target gene. We targeted the *OCT4* gene, given the inconsistency of results from previous reports. We hypothesized that ribonucleoproteins formed with CRISPR-Cas9D10A produce larger deletions at greater consistency and efficiency than CRISPR-Cas9, and CRISPR-Cas13a can efficiently knockdown mRNA in cattle zygotes. We also hypothesized that simultaneous targeting of DNA and RNA could ablate gene function in cattle zygotes *in vitro*. In this study, we have mitigated the barriers of poor deletion efficiency and the presence of preexisting mRNA while maintaining embryo survival. The dual delivery of CRISPR-Cas9D10A, six-hours apart, increases the incidence of gene editing and full deletions. Additionally, we targeted maternally inherited transcripts with CRISPR-Cas13a while simultaneously removing a targeted sequence of the genome. Altogether, we have developed a method for high efficiency genome and transcriptome editing in bovine zygotes using CRISPR-Cas editing technology. Our approach overcomes many limitations of gene editing for mechanistic studies of gene function in pre-implantation embryos. Although cattle blastocyst formation is possible in the absence of OCT4, these embryos lack an inner cell mass and present severe transcriptional dysregulation of several genes related to stemness.

## Results

First, we assessed the efficacy of electroporation and the cleavage function of the ribonucleoproteins (RNPs). Here, we used electroporation conditions modified from a previous publication (35), as follows: six poring pulses of 15 volts, with 10% decay, for two milliseconds with a 50-millisecond interval, immediately followed by five transfer pulses of 3 volts, 40% decay, for 50 milliseconds with a 50-millisecond interval, alternating the polarity. Fluorescence imaging showed that the RNP formed by Cas9-RFP + scramble guide RNAs (gRNAs) bypassed the zona pellucida in nearly all putative zygotes (PZ) electroporated (Fig. 1A). Next, we confirmed that the RNPs formed with Cas9 + OCT4 single guide RNA (sgRNA) 1 or Cas9 + OCT4sgRNA2 were able to cleave the targeted DNA *in vitro* (Fig. 1B).

**Fig. 1.**
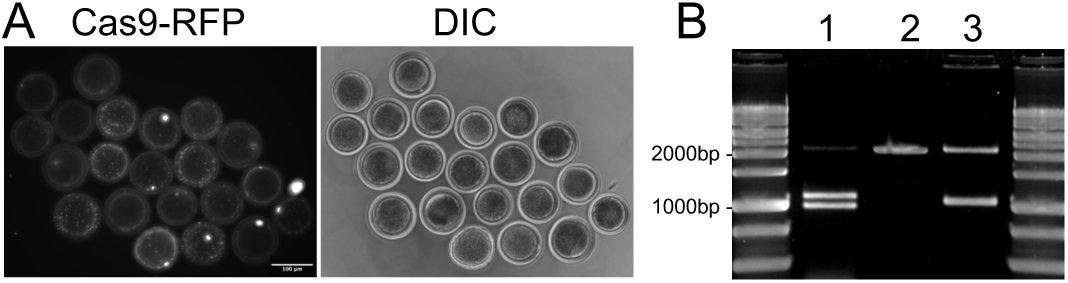
CRISPR-Cas in cattle zygotes. (A) Putative zygotes electroporated with Cas9-RFP. (B) *In vitro* cleavage assay showing the targeted cleavage of DNA by CRISPR-Cas9 and specific sgRNAs. 1: targeted DNA fragment treated with Cas9+sgRNA1, 2: uncut, original DNA fragment; 3: targeted DNA fragment treated with Cas9+sgRNA2.

### Both CRISPR-Cas9 and CRISPR-Cas9D10A produce deletions in cattle zygotes

First, we asked if Cas9 and Cas9D10A would result in similar editing efficiencies and deletion patterns. High-throughput targeted sequencing revealed that 73.1% and 81.5% of embryos presented at least one segment of DNA deleted when we used Cas9 (N embryos = 26) or Cas9D10A (N embryos = 27), respectively. We observed that 15.4% and 25.9% of the embryos electroporated with Cas9 or Cas9D10A, respectively, and genotyped by sequencing, did not have a wild-type copy of the DNA in the targeted region. The deletions resultant from Cas9 or Cas9D10A varied in their location and length. We observed that Cas9D10A RNPs produced longer deletions and removed the segment of DNA that included both sgRNAs, whereas Cas9 mostly produced small deletions in the region surrounding the sgRNAs but did not cause many deletions spanning both sgRNAs.

Although not significant (P=0.27, Fisher’s Exact test), Cas9D10A produced 10.5 percentage points more full deletions, with no wildtype alleles, when compared to Cas9. Thus, we carried out the next experiments with Cas9D10A and *OCT4*-targeting sgRNAs. Also, considering that many of the Cas9-RFP RNPs remained in the membrane or perivitelline space (Fig. 1A), we reasoned that a second electroporation would increase the efficiency of full deletions in cattle presumptive zygotes (PZ). A second electroporation of PZ (approximately six hours after the first electroporation; see methods for details) with RNPs composed of Cas9D10A and associated sgRNAs resulted in no PCR amplification for most blastocysts when using the oligonucleotides designed for high-throughput short-read sequencing (Fig. S1A, B, Appendix). This outcome, and prior reports that CRISPR-Cas9 can produce unexpected large deletions (60, 61), prompted us to design oligonucleotides to flank a wider region of the DNA surrounding our sgRNA target sequences. Approximately 19% of the blastocysts tested with this long-range pair of oligonucleotides produced an amplicon (Fig. S1C, Appendix). All blastocysts that had no amplicon produced with oligos surrounding our targeting sgRNAs were tested for amplification of a non-targeted autosomal region of the genome to confirm that an embryo was present in the tube (Fig. S1D, E, Appendix).

We sequenced the PCR products from seven blastocysts using the Sanger procedure, and three of these samples produced electropherograms from only one fragment (Fig. 3A). The long-range PCR produced multiple amplicons in the other three samples, which is unsuitable for Sanger sequencing. Therefore, we decided to proceed with Nanopore sequencing for multiple allele detection. Twenty-four blastocysts produced amplicons with long-range oligos and were sequenced by Sanger (Fig. 3A) or Nanopore (Fig. 3B) methods. Sequencing results showed that 95.8% (23/24) of the blastocysts had at least one chromosome with a deleted segment on the targeted DNA sequence (see an example in Fig. S2, Appendix). In addition, 70% (17/24) of the blastocysts sequenced did not present a wild-type sequence in the targeted region (Fig. 3B, other two examples in Fig. S3, Appendix). We note that 72 out of 89 blastocysts tested with our long-range oligos did not produce an amplicon, though the presence of DNA was confirmed by amplification of a non-targeted autosomal region in each sample. Therefore, we can reason that these 72 blastocysts had unexpectedly larger DNA deletions (60, 61) that eliminated at least one oligonucleotide pairing site on all chromosomes. Under such reasoning, we can estimate that 91% (81/89) of the blastocysts were edited with no wild-type sequence of the targeted DNA.

**Fig. 2.**
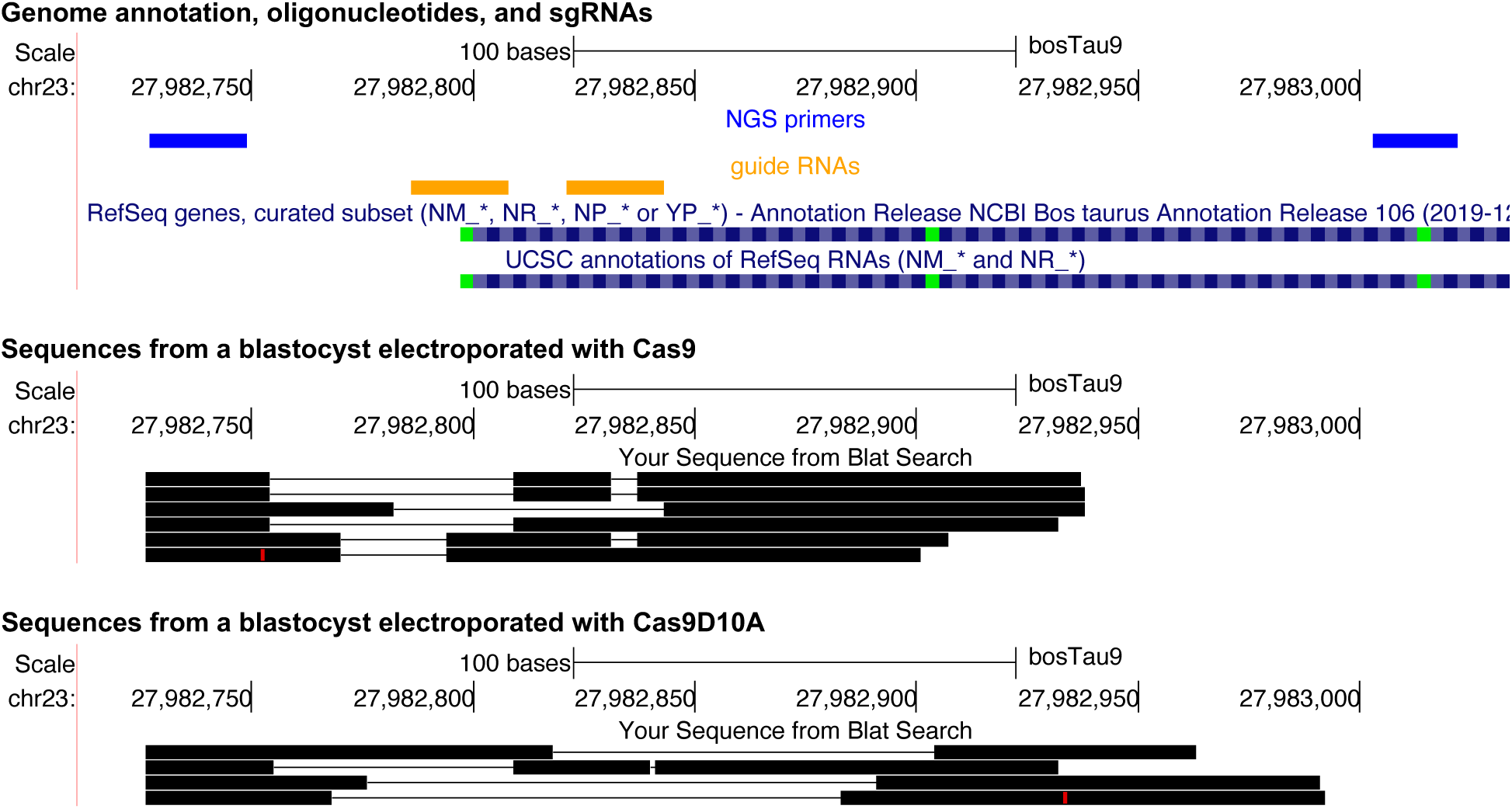
Targeted DNA deletions using CRISPR-Cas9. Representative images of the DNA mapping of sequences resultant from high-throughput sequencing of embryos electroporated with either Cas9 or Cas9D10A aligned to the cattle genome.

**Fig. 3.**
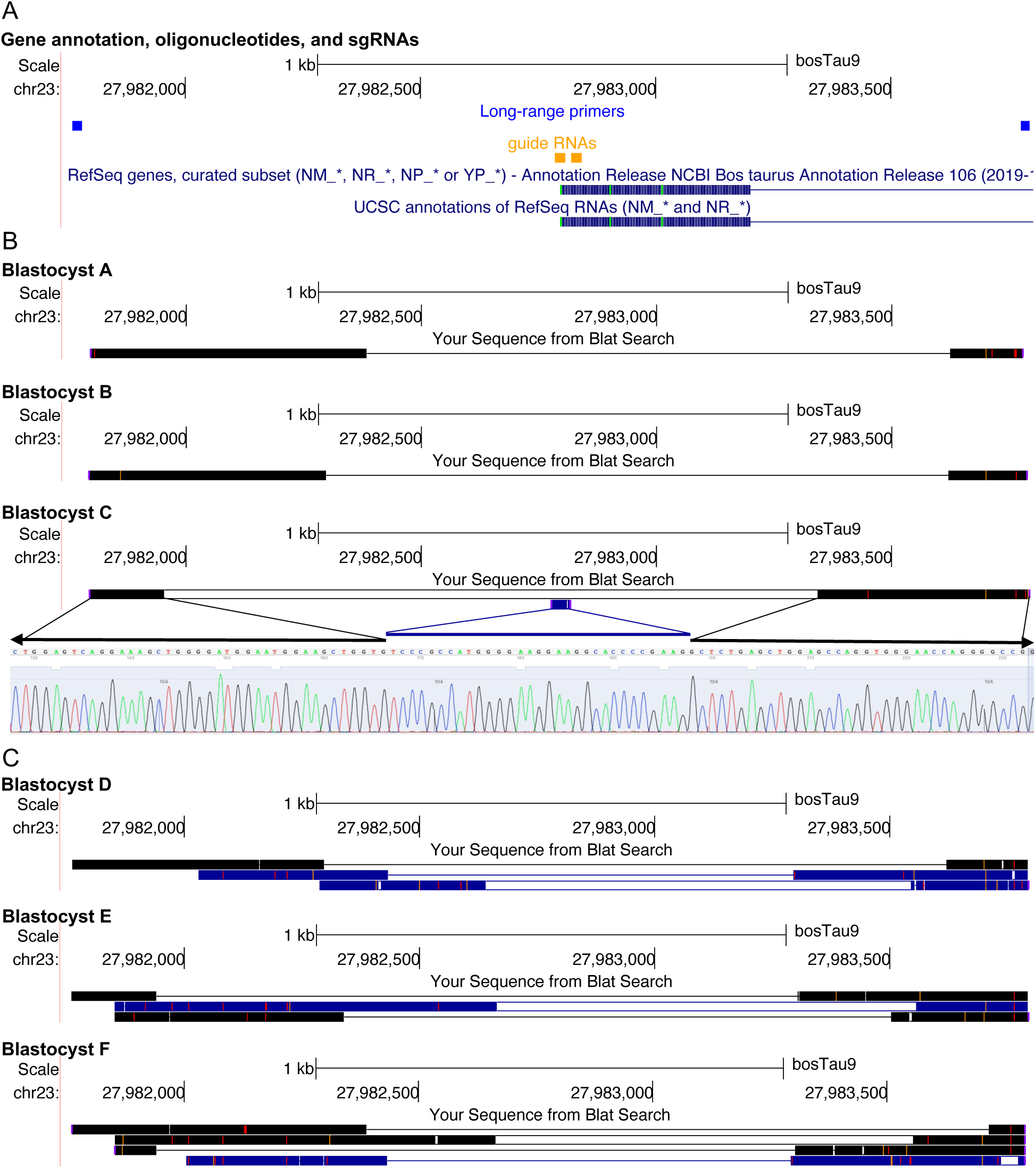
Representative schematics of DNA mapping from fully edited blastocysts produced by two sessions of electroporation with Cas9D10A and *OCT4*-targeting sgRNAs. (A) Genome annotation identifying sgRNA targets and sequencing primers. (B) Sanger and (B) Nanopore targeted sequencing.

### Embryo survival following one or two Cas9D10A electroporation sessions

We tested if the electroporation of Cas9D10A with scramble gRNAs would impact development to the blastocyst stage. One electroporation session with scramble gRNAs produced similar results to controls (164-166 hpf – Cas9D10A and scramble gRNAs: 17.1%±3.1, controls: 25.3%±3.2; 188-190 hpf – Cas9D10A and scramble gRNAs: 31.5%±3.8, controls: 30.8%±3.4, P>0.05, Tables S1-3, Appendix). Two electroporation sessions with scramble gRNAs also produced similar results to controls (164-166 hpf – Cas9D10A and scramble gRNAs: 17.7%±2.6, controls: 25.3%±3.2; 188-190 hpf – Cas9D10A and scramble gRNAs: 28.2±3.1, controls: 30.8%±3.4, P>0.05, Tables S1-3, Appendix). Therefore, one or two electroporation sessions with Cas9D10A and scramble gRNAs did not reduce blastocyst yield and maintained survival like that seen in non-electroporated embryos.

One electroporation session with Cas9D10A and *OCT4*-targeting sgRNAs reduced the blastocyst yield relative to scramble or control groups (164-166 hpf – Cas9D10A and targeting sgRNAs: 6.8%±1.4, controls: 25.3%±3.2; 188-190 hpf – Cas9D10A and targeting sgRNAs: 11.6%±1.7, controls: 30.8%±3.4, P<0.001, Tables S1-3, Appendix). Two electroporation sessions with Cas9D10A and *OCT4*-targeting sgRNAs also reduced blastocyst development (164-166 hpf – Cas9D10A and targeting sgRNAs: 3.2%±0.7, controls: 25.3%±3.2; 188-190 hpf – Cas9D10A and targeting sgRNAs: 7.9%±1.1, controls: 30.8%±3.4, P<0.001, Tables S1-3, Appendix). We also evaluated zygotes electroporated twice with Cas9D10A and *OCT4*-targeting sgRNAs (N=56), transferred into individual drops of media at the 8-cell stage and placed in a time-lapse incubator, along with controls that were not electroporated (N=28). A greater number of electroporated embryos arrested their development at the 8-cell (35.5% vs 17.8% controls, P=0.0013) and morula (51.8% vs 35.7% controls, P=0.0013) stages. Additionally, a lower proportion of the electroporated embryos developed to the blastocyst stage (12.5% vs 46.4% controls, P=1.06×10^−7^, exact binomial test). Thus, targeting the gene *OCT4* by two electroporation sessions of Cas9D10A and sgRNAs caused partial developmental arrest at the 8-cell and morula stages with a sharp decline in the development to the blastocyst stage but did not eliminate embryo survival.

### mRNA knockdown in cattle zygotes by Cas13a

To test whether Cas13a can target mRNAs in zygotes, first, we electroporated PZ with exogenous mRNAs of fluorescent proteins (red (RFP) or green (GFP)). Fluorescence imaging of embryos ∼70 hpf showed successful introgression of exogenous mRNAs (GFP and RFP mRNAs) into PZ and expression of the corresponding proteins in cleavage embryos (Fig. 4). By contrast, we quantified a significant reduction of fluorescence (1.37-fold for GFP, and 1.34-fold for RFP, P<0.001, Fig. 4) when we electroporated PZ with the exogenous mRNA and Cas13a + targeting sgRNAs simultaneously. Since those PZ treated with Cas13a + targeting sgRNAs did not target an endogenous RNA, we tested whether Cas13a RNPs would impact embryo development. There were no statistical differences in the development to blastocyst stage at 188-190 hpf (30.8%, 37.7%, 32.1%, 33.3% for Cas13a+GFP mRNA sgRNAs, Cas13a+RFP mRNA sgRNAs, Cas13a+scramble gRNAs, controls, respectively, P>0.8). Thus, Cas13a targets mRNAs efficiently in cattle zygotes with no alteration in their developmental potential.

**Fig. 4.**
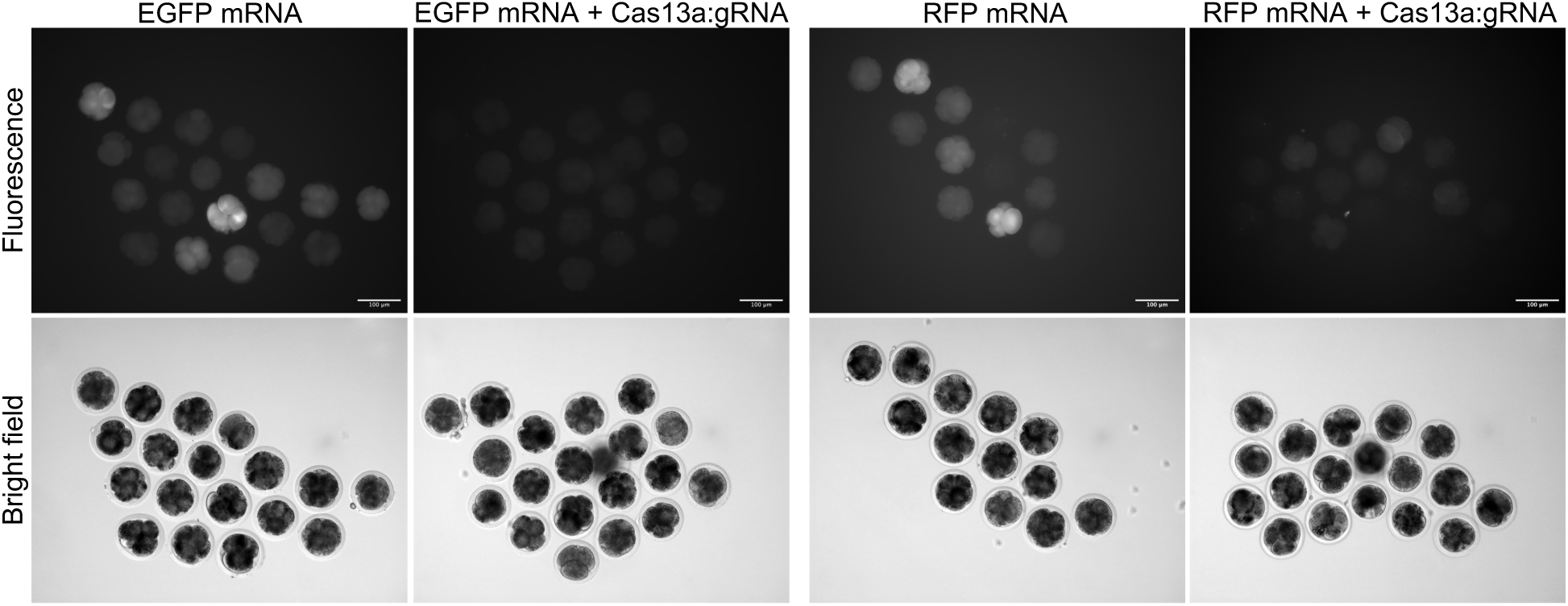
Knockdown activity of Cas13a in cleavage cattle embryos. Scale bar: 100μm.

### Ablation of OCT4 function in cattle pre-implantation embryos by CRISPR-DART

We used CRISPR-DART (Fig. 5) to target the promoter (based on orthology with the human genome) and exon 1 of *OCT4*. The induced deletions significantly reduced embryo survival (164-166 hpf – CRISPR-DART: 6.1%±0.8, controls: 23%±2.3; 188-190 hpf – CRISPR-DART: 13%±1.2, controls: 31.6%±2.5, P<0.001, Tables S4-S5, Appendix). Using immunofluorescent staining, we determined that the putatively edited blastocysts (we estimated 91% editing success) did not produce OCT4 protein. Additionally, we detected a decrease of NANOG in the edited blastocysts (Fig. 6A). Thus, we confirmed that the deletion of the promoter and exon 1 of *OCT4* resulted in absence of OCT4 protein.

**Fig. 5.**
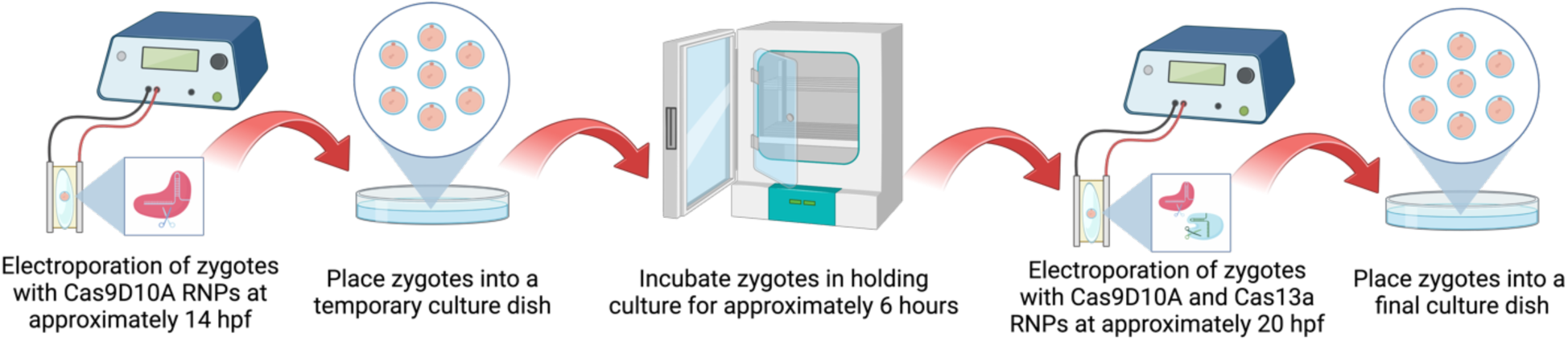
Schematic of CRISPR-DART procedure. Created with BioRender.com.

**Fig. 6.**
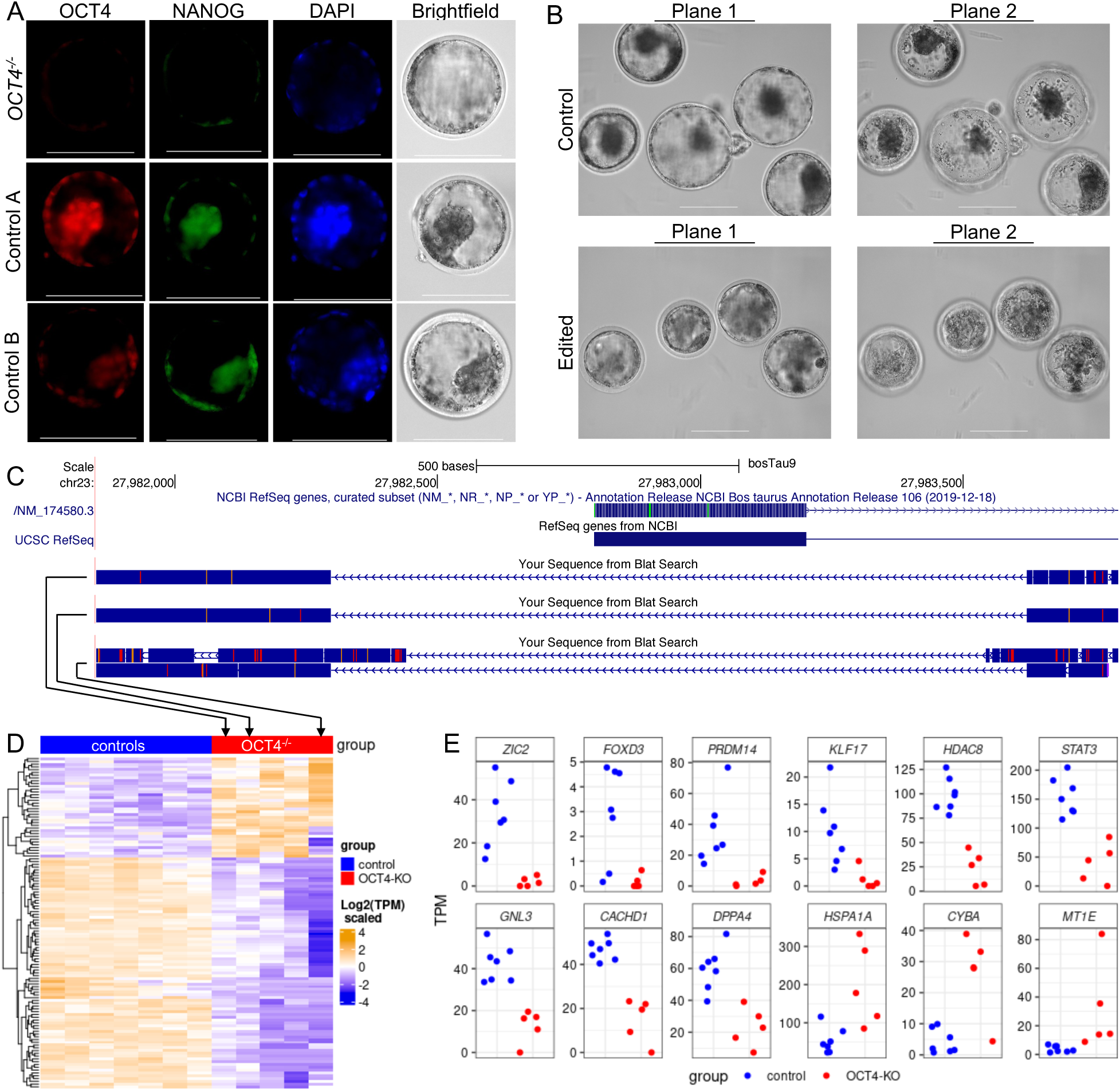
Impact of *OCT4* knockout in cattle pre-implantation embryos. (A) Immunofluorescence assay of OCT4 and NANOG in cattle pre-implantation embryos. Scale bar: 100 µm. (B) *In vitro* produced blastocysts 188-190 hpf. Images are presented in two focal planes for the visualization of the inner cell mass and blastocoel cavity. Scale bar: 100 µm. (C) Schematics of the DNA sequence mapping from three of five blastocysts used for RNA-sequencing. (D) Heatmap depicting the relative differential transcript abundance of 125 genes in *OCT4* knockout blastocysts. (E) Transcript abundance of 12 genes functionally associated with the maintenance of pluripotency.

Morphological examination showed an absence of a well-defined inner cell mass in blastocysts deemed *OCT4*^−/−^, whereas a well-defined inner cell mass is clearly visible in the control embryos (Fig. 6B). Time-lapse image analysis of the development of putative *OCT4*^−/−^ embryos confirmed the formation of a blastocoel cavity and absence of a normal inner cell mass (Supplementary movies 1 and 2). Next, we interrogated the transcriptome of *OCT4*^−/−^ blastocysts. To confirm that the blastocysts collected were *OCT4*^−/−^, we obtained genomic DNA and total RNA from single embryos. We used the genomic DNA to confirm the absence of the targeted DNA sequence (Fig. 6C) and evaluated the transcript abundance of 14156 protein-coding or long non-coding genes from five *OCT4*^−/−^ blastocysts. Comparative analyses revealed 125 genes with differential transcript abundance between *OCT4*^−/−^ blastocysts and controls (Fig. 6D, FDR<0.1). Eighty-three and 42 genes had lower and greater transcript abundance in *OCT4*^−/−^ blastocysts, respectively. Notably, 17 genes with differential transcript abundance were functionally related to the maintenance of pluripotency (see Fig. 6E for examples). These results indicate that a blastocoel cavity may form in the absence of *OCT4*, but the formation of the inner cell mass and normal gene expression is severely impaired in cattle *OCT4*^−/−^ embryos.

## Discussion

We developed an approach using Cas9D10A to delete targeted regions of the DNA and Cas13a to cleave targeted RNA for complete disruption of gene function in cattle zygotes at high efficiency. We used CRISPR-DART to target *OCT4* mRNAs and exon 1, including the promoter region. Our data provide several insights into the function of *OCT4* in cattle pre-implantation development. First, most *OCT4*^−/−^ embryos arrest development before the blastocyst stage, but a minor proportion of edited zygotes do still survive. Second, *OCT4*^−/−^ embryos that progress their development are able to form a blastocoel cavity with an outer layer of cells resembling trophectoderm but do not form an inner cell mass with similar morphology observed in control embryos. Finally, the ablation of *OCT4* significantly alters the transcript abundance of genes involved in pluripotency. Our results show that *OCT4* is necessary for the development of a cattle blastocyst with a morphologically normal inner cell mass.

### Simultaneous deletion of DNA segments and cleavage of RNA in zygotes

Previous research has reported the use of CRISPR-Cas9 to delete DNA segments in cattle zygotes (32, 35–37, 40). To build on this, we tested the efficacy of Cas9 and Cas9D10A with two sgRNAs targeting the exon 1 of *OCT4*. Although we did not test for off-targets, Cas9D10A produces single-strand DNA breaks and requires two sgRNAs targeting opposite strands to nick the DNA and induce faulty DNA repair (62). This combination of factors significantly reduces mutation elsewhere in the genome. Our results confirmed that Cas9D10A RNPs produce large deletions beyond the region flanked by the sgRNAs (63). However, we only detected deletions larger than 500 nucleotides when we electroporated the zygotes twice in an interval of six-hours between sessions. Two electroporation sessions allow for the introduction of greater quantities of RNPs in the zygote without causing toxicity (64). The combination of Cas9D10A targeting two sequences in the genome, and likely a higher quantity of RNPs entering the cell in two sessions of electroporation, increased the efficiency in producing full edits from 25.9 to 91%, which is higher than previous reports in cattle zygotes (32, 35–37, 40).

The RNPs produced by the combination of CRISPR-Cas13a and an sgRNA can target and cleave single stranded RNAs (65, 66). These RNPs have been used in animal embryos including in mouse (67) and pig (67) to target specific mRNAs. Here, we tested the efficacy of Cas13a in cattle zygotes by introducing and targeting mRNAs for either GFP or RFP. Our experiments showed that Cas13a could efficiently prevent protein synthesis from targeted mRNAs in cattle zygotes. One concern related to Cas13a is that it may cleave unintended mRNAs in the vicinities of targeted RNAs in a cell-dependent manner (68). The introduction of Cas13a+sgRNAs targeting exogenous mRNAs (GFP or RFP) and the corresponding mRNAs into cattle zygotes did not reduce embryo survival, thus, if there are off-target effects, they are negligible in cattle pre-implantation embryos. Cas13a can knockdown specific mRNAs in zygotes in conjunction with Cas9D10A to target genomic DNA.

### Effects of ablation of OCT4 in cattle embryos

Our CRISPR-DART approach efficiently deleted exon 1 and the promoter from the *OCT4* in most embryos. We expected that removing the promoter and transcript starting site would impair the production of *OCT4* mRNAs and proteins. Indeed, we confirmed that most of the embryos tested by immunofluorescence assays did not have detectable OCT4. However, our RNA-sequencing data, produced from confirmed *OCT4*^−/−^ embryos, showed sequences aligning with all five exons of *OCT4* (Fig. S4A, Appendix). Both data are conflicting, but we reason that a pseudogene of *OCT4* produced RNAs that were sequenced and mapped to the *OCT4* gene. Few lines of evidence support our rationale. We only selected sequences that mapped to the cattle reference genome once to quantify transcript abundance. Messenger RNAs produced by this pseudogene would have mapped to both genomic regions, but this region is not present in the current cattle reference genome, as indicated by the comparative mapping in the UCSC genome browser (Fig. S4B, Appendix). Annotated *OCT4* pseudogenes are long non-coding RNAs (69–74) and do not contain introns (Fig. S4B, Appendix). Lastly, one hallmark observation in our data is that no sequence mapped to *OCT4* intronic regions in edited embryos (Fig. S4A, Appendix), whereas several intronic sequences from all introns were evident in the control embryos (Fig. S4C, Appendix). Thus, we concluded that the RNA sequences mapping to the annotated *OCT4* cattle gene for *OCT4*^−/−^ embryos were transcribed from a pseudogene with no intron. This confounding factor between functional *OCT4* and pseudogenes has been long observed in stem cell research (71).

The ablation of *OCT4* function in cattle pre-implantation embryos severely reduced blastocyst development, but a majority of the blastocysts were confirmed to be fully edited. This finding aligns with reports that produced embryos from *OCT4^−/−^* somatic cells (56, 57) or produced putative *OCT4^−/−^* embryos by introducing RNPs into zygotes (35, 36). The major blastocyst phenotype we observed was the absence of an inner cell mass, a phenotype previously reported in knockout mice (Pou5f1^tm1Cgre^/Pou5f1^tm1Cgre^, genotype id MGI:3040797 (49)). By comparison, Simmet and colleagues showed that *OCT4* is necessary only for the second lineage differentiation (57), and it is possible that dysregulated genomic reprogramming due to somatic cell cloning could be the cause of the minor discrepancy in the phenotypes. Our results show that *OCT4* is required for the differentiation of inner cell mass in cattle embryos.

We also evaluated the transcriptomic profile of *OCT4^−/−^*blastocysts. Our results, contrasting the transcriptome of *OCT4^−/−^ in vitro* produced blastocysts and controls, showed 125 genes with significant alteration in transcript abundance. Only seven genes overlapped with the dataset reported elsewhere (*ARF5*, *MT1E*, *NUPR1*, *PLA2G15*, *RRM2*, *STAT3*, and *SWAP70* (57)), but all seven genes had the same direction of altered transcript abundance. Among the genes up-regulated in *OCT4^−/−^* blastocysts, *MT1E* potentiates cell differentiation (75, 76), and *NUPR1* is activated when the cells are under oxidative stress (77). Conversely, among the genes with lower transcript abundance in *OCT4^−/−^*blastocysts, *RRM2* is a known marker of pluripotent stem cells (78), *STAT3* is required for embryonic stem cell pluripotency (79), and the absence of *SWAP70* impairs the self-renewal of mouse hematopoietic stem cells (80).

Our results also highlighted a dysregulation in other genes with roles in regulating stem cell function. For example, *DPPA4* and *CADHD1* are pluripotent stem cell markers (78) and have a lower abundance of transcripts in *OCT4^−/−^* blastocysts. Also, with a lower abundance of transcripts in edited embryos, *ZIC2* (81, 82), *KLF17* (83, 84), *FOXD3* (85), *HDAC8* (85), and *GNL3* (86) are directly involved in the regulation of pluripotency. By contrast, *ADAM9* (87), *ANXA8L1* (87), *CDKN1A* (88), *DDAH2* (89), *ELF3* (89), *HOXC10* (90) and *LGI1* (90) are all up-regulated in *OCT4^−/−^*blastocysts and promote cellular differentiation. Collectively, these results show a severe imbalance in the regulation of genes associated with stemness or cell differentiation, which is coherent with the absence of an inner cell mass in *OCT4^−/−^* blastocysts.

### Limitations of the study

This study has limitations, mostly related to the editing efficiency and the inherent biology of DNA synthesis in zygotes. First, several technical factors, such as the concentration of ribonucleoprotein and electroporation conditions, may be finetuned to improve the efficiency of producing fully edited embryos. Here, we based our technical conditions on previous literature (34, 35), but we are confident there is room for improvement. Second, although matured oocytes are exposed to sperm in the fertilization media simultaneously, there is a window of opportunity for oocytes to be fertilized. Thus, among hundreds of oocytes, the timing of fertilization is heterogeneous. Third, following fertilization, both pronuclei are formed of compacted chromatin (91, 92), in which the RNPs are likely not accessible to the targeted sequence. The unwinding of the DNA for synthesis is an asynchronous process across zygotes, and DNA synthesis can happen in a window of approximately 10 hours (32, 93). Therefore, there is tremendous variability in the accessibility of the targeted DNA across zygotes. Fifth, we did not sequence the DNA of embryos that had deletions larger than the DNA sequence flanked by our oligonucleotides. We made several attempts to amplify very large fragments of 6 and 8 kilobases long, but nonspecific amplification hindered our ability to genotype the embryos accurately. Lastly, we had a relatively small sample size for the transcriptome analysis (five *OCT4^−/−^* and seven control blastocysts), but we countered this limitation with stringent analysis and scrutiny of the results based on the literature. Despite these limitations, our results show improvement in the efficiency of producing fully edited embryos, and the phenotype observed is coherent with the literature in mice and cattle.

## Conclusion

The production of knockouts is essential for mechanistic studies of gene function in pre-implantation embryos. We showed that Cas9D10A is more efficient than Cas9 at producing biallelic deletions in zygotes. Two sessions of electroporation introduce greater quantities of Cas9D10A RNPs and increase the frequency of large biallelic deletions. The sequential introduction of RNPs does not impair embryo development as long sgRNAs targeting proximal sequences in the genome are not used. RNPs consisting of Cas13a prevent protein production from targeted mRNAs in cattle zygotes. Our CRISPR-DART approach increased the efficiency of producing knockout zygotes. Lastly, we show that *OCT4* is required for the regulation of several genes that control pluripotency and the formation of an inner cell mass in cattle blastocysts.

## Materials and Methods

### In vitro production of embryos

Unless otherwise specified, all reagents were purchased from Sigma-Aldrich.

All procedures and culture media composition for *in vitro* production of embryos are described in detail elsewhere (94, 95). Briefly, we obtained cattle ovaries from an abattoir (Brown Packing, Gaffney, SC) and washed them with anti-biotic anti-mycotic (Antibiotic-Antimycotic 100X, Thermofisher Scientific, Waltham, MA) and 0.9% saline solution. For the collection of cumulus-oocyte-complexes (COCs), we aspirated ovarian follicles 3-8mm in diameter using an 18g needle (Single-Use Needles BD Medical, VWR, Philadelphia, PA) connected to a regulated vacuum system and collection bottle containing oocyte collection medium (OCM, BoviPlus Oocyte Collection Medium, Minitube, Verona, WI) supplemented with gentamicin (50 µg/µl) and heparin (2 U/ml). We washed COCs twice in OCM, followed by three washes in oocyte maturation medium (OMM). Then we selected COCs with homogeneous, non-granular oocyte cytoplasm and three or more compact layers of cumulus for *in vitro* maturation. COCs were placed in groups of 10 in 50 µl of OMM covered by light mineral oil. *In vitro* maturation plates were incubated for 22-24 hours at 38.5°C and 5% CO_2_ humidified atmosphere. Following the incubation, we washed the mature COCs in synthetic oviductal fluid medium (SOF) containing N-2-hydroxyethylpiperazine-N’-2-ethanesulfonic acid (HEPES-TL, Thermofisher Scientific, Waltham, MA) and SOF for fertilization (SOF-FERT) before transferring into a final fertilization plate (100 COCs/ml). We thawed frozen semen straws and processed sperm prior to transfer into fertilization plates at a concentration of 1,000,000 spermatozoa/ml. COCs and spermatozoa were co-incubated for 12-13 hours under the same conditions described for *in vitro* maturation.

We removed putative zygotes (PZ) from fertilization medium at approximately 14 hours post fertilization (hpf) and denuded the cumulus cells by vortexing in 1% hyaluronidase for 5 minutes. Next, we moved PZ through three washes of SOF-HEPES and SOF culture medium (SOF-BE1). The PZs used for control groups were placed in their final culture dish immediately after the washes. Alternatively, the PZs used for electroporation were placed in temporary culture dishes containing 50µl SOF-BE1 covered with light mineral oil. After electroporation, we washed the PZs in SOF-BE1 before placing them in culture. PZs were cultured in groups of 25-30 in 50 µl SOF-BE1 covered by light mineral oil, incubated at 38.5°C with 5% CO_2_, 5% O_2_ in a humidified Eve Benchtop Incubator (WTA, College Station, TX).

For time-lapse image analysis, we cultured 8-cell embryos individually in 15 µl SOF-BE1 covered by light mineral oil, incubated at 38.5°C with 5% CO_2_ and 5% O_2_ in a MIRI Time-Lapse Incubator (Esco Medical, Egaa, DK).

### Guide RNA design

We designed sgRNAs to target the genomic DNA of the transcriptional start site and exon 1 of *OCT4* using the CRISPOR webservice (96). We designed the sgRNAs for Cas13a using New York Genome’s cas13designtool (97, 98) to target the 4th exon of the *OCT4* mRNA. As an independent layer of *in silico* validation, we aligned all sgRNAs targeting the *OCT4* gene or transcript to the bovine genome with the BLAT software in the UCSC Genome Browser (99). Additionally, Cas13a sgRNAs were designed to target CleanCap EGFP and mCherry mRNAs (5moU, TriLink Biotechnologies, San Diego, CA). The targeting sgRNAs used in this study were OCT4 sgRNA1: CTTCGCCTTCTCGCCCCCGCCGG, OCT4 sgRNA2: TGTCCCGCCATGGGGAAGGAAGG, OCT4 mRNA sgRNA: ATGCTCTCCAGGTTGCCTCT, mCherry mRNA sgRNA: TCCTCGAAGTTCATCACCCG, EGFP mRNA sgRNA: CATGATATAGACGTTGTGG. We purchased all sgRNAs as a single RNA molecule comprised of both crRNA and tracrRNA sequences (Integrated DNA Technologies (IDT), Research Triangle Park, NC). We also purchased a scramble gRNA (Alt-R® CRISPR-Cas9 Negative Control crRNA #1) and tracrRNA (Alt-R® CRISPR-Cas9 tracrRNA) from IDT and combined them following the manufacturer’s instructions.

### Preparation of ribonucleoprotein and procedures for electroporation

We mixed Cas9 and sgRNAs for the formation of ribonucleoproteins in Opti-MEM™ Reduced Serum Medium (Thermofisher Scientific, Waltham, MA), and maintained the solution at room temperature for at least 30 minutes prior to electroporation. The specific concentrations and enzymes are detailed below.

As detailed above for control cultures, we removed the cumulus cells from the PZ and placed them in holding SOF-BE1 at 38.5°C, 5% CO_2_, and 5% O_2_. We removed PZs in groups of 30-40 from a holding culture and briefly washed them in OptiMEM (previously equilibrated in the incubator at 38.5°C and 5% CO_2_). Next, we mixed 3 µl of the solution containing ribonucleoproteins with 3 µl of OptiMEM containing PZs. We carried out the electroporation using a BTX oocyte petri dish with platinum electrodes (Harvard Apparatus, VWR, Philadelphia, PA). We transferred the final 6 µl to the electroporation chamber. Impedance was checked and, if necessary, adjusted to measure between 0.19 and 0.20 by the addition of OptiMEM or removal of the electroporation solution. The electroporation parameters were as follows: six poring pulses of 15 volts, with 10% decay, for two milliseconds with a 50-millisecond interval, immediately followed by five transfer pulses of 3 volts, 40% decay, for 50 milliseconds with a 50-millisecond interval, alternating the polarity. Following the electroporation, we washed the PZ with OptiMEM and SOF-BE1.

### Cleavage assay of the targeted DNA

We carried out a cleavage assay to assess the formation and cleavage of DNA by ribonucleoproteins (100). We amplified a segment of genomic cattle DNA to be targeted by the sgRNAs by assaying a PCR using the following oligonucleotides (forward: GGCAAGGAACTTGATGCACG and reverse: TGGCCAACCCACTGTTTGAT). The PCR reaction mix consisted of 0.2 IU/μl Phusion Hot Start II DNA Polymerase (Thermofisher Scientific, Waltham, MA), 1X Phusion HF Buffer, 200 μM dNTPs (Promega, Madison, WI), and forward and reverse oligonucleotides (IDT, Coralville, Iowa) at 0.10 μM each, in a final volume of 20 μl in 0.2 ml clear PCR tubes. The cycling conditions for this reaction were: 98°C for 1 minute, followed by 40 cycles of 98°C for 15 seconds, 55°C for 45 seconds, and 72°C for 1 minute, followed by a final extension of 4 minutes at 72°C.

We incubated Cas9 (1 μM, Integrated DNA Technologies, Research Triangle Park, NC) with either sgRNA1 or sgRNA2 300 nM in OptiMEM for 30 minutes at room temperature to form the RNPs. Next, we incubated RNPs with DNA fragments containing the targeted sequence (1:10 (v:v) Cas9+sgRNA, 3 nM DNA, 1x NEB buffer 3.1) at 37°C for 3 hours. Fragments were assessed by electrophoresis on a 1.5% Agarose I™ gel followed by staining with Diamond™ Nucleic Acid Dye and imaging.

### Evaluation of electroporation efficiency

We evaluated the electroporation efficiency with RNPs formed by Cas9-RFP (Alt-R™ S.p.Cas9-RFP V3, Integrated DNA Technologies, Research Triangle Park, NC) at 800ng/µl and scramble gRNAs at 800ng/µl. After washing the PZs in OptiMEM, we imaged them using a fluorescent microscope (details below).

### Assessment of sequence deletions by Cas9 or Cas9D10A

To test the pattern of deletions with either a double-cutting enzyme or a nickase, we carried out a single electroporation at approximately 15 hpf with ribonucleoproteins formed by either Cas9 or Cas9D10A (IDT) at 800ng/µl and sgRNAs at 800 ng/µl each. After washing the PZ in SOF-BE1, we placed them in culture as described for control PZs.

### Assessment of mRNA cleavage by Cas13a

We carried out a single electroporation of PZs with one of the following solutions: a) mRNA of either mCherry or GFP at 400ng/µl; or b) mRNA of either mCherry or GFP at 400ng/µl and ribonucleoprotein formed by Cas13a (GenScript, Piscataway, NJ) at 400 ng/µl and the corresponding targeting sgRNA at 400 ng/µl. After washing the PZ in OptiMEM, we imaged them using a fluorescent microscope (details below) in SOF-HEPES.

### CRISPR-DART

For CRISPR-DART, we carried out the first electroporation at approximately 14 hpf with 3µl of RNPs formed by Cas9D10A at 600ng/µl and sgRNAs at 800 ng/µl each mixed with 3µl of OptiMEM. The PZs were maintained in SOF-BE1 media in the incubator. Then, we electroporated them again at approximately 20 hpf with two solutions of RNP complexes prepared separately. One solution contained Cas9D10A at 600ng/µl and each sgRNA at 800 ng/µl and the other contained Cas13a at 1600 ng/µl and sgRNA at 800 ng/µl. At the time of electroporation, we mixed 1.5 µl of each RNP with 3 µl of OptiMEM containing the PZ. After washing the PZ in SOF-BE1, we placed them in culture as described for control PZs.

### Targeted DNA sequencing

All embryos collected for DNA sequencing were washed in PBS 0.1% BSA fraction V, followed by removal of the zona pellucida by exposure to EmbryoMax® Acidic Tyrode′s Solution and gentle pipetting. Once the zona pellucida was removed, we washed the embryos in PBS 0.1% BSA fraction V twice and collected them individually in microtubes in approximately 1 µl PBS 0.1% BSA fraction V. We exposed the nucleic acids of each embryo with 5µl of QuickExtract™ DNA Extraction Solution (Lucigen, VWR, Philadelphia, PA), and incubated at 65°C for 15 minutes followed by 2 minutes at 98°C.

### High throughput short reads

We assayed a PCR using oligonucleotides flanking the targeted deletion site with coupled sequencing adapters on their 5’ end (forward: acactctttccctacacgacgctcttccgatctAGAGGTGTTGAGCAGTCTCTAGG, reverse: gtgactggagttcagacgtgtgctcttccgatctGTAGGCCATCCCTCCACAC; lower case letters indicate adapter, uppercase letters indicate targeted sequence). The PCR reaction consisted of 0.2 IU/μl Phusion Hot Start II DNA Polymerase (Thermofisher Scientific, Waltham, MA), 1X Phusion HF Buffer, 200 μM dNTPs (Promega, Madison, WI), and oligonucleotides (IDT) at 0.1 μM each, in a final volume of 20 μl. Reactions were carried out in 0.2 ml clear PCR tubes (VWR, Philadelphia, PA), and the cycling conditions were: 98°C for 1 minute, followed by 40 cycles of 98°C for 15 seconds, 61°C for 30 seconds, and 72°C for 40 seconds, proceeding a final extension of 4 minutes at 72°C. We confirmed the amplification using 2% Agarose I™ (VWR, Philadelphia, PA) and gel electrophoresis, followed by DNA staining with Diamond™ Nucleic Acid Dye (Promega, Madison, WI)

Next, we completed the library preparation with a second PCR using oligonucleotides obtained from xGen™ UDI Primers Plate 1, 8nt (IDT). The reaction mixture consisted of 0.3 IU/μl Phusion Hot Start II DNA Polymerase (Thermofisher Scientific, Waltham, MA), 1X Phusion HF Buffer, 200 μM dNTPs (Promega, Madison, WI), and 3 µM Illumina adaptors, in a final volume of 25 μl. The reaction was assayed according to the following conditions: 98°C for 30 seconds, followed by 15 cycles of 98°C for 10 seconds, 60°C for 30 seconds, and 72°C for 30 seconds, proceeding a final extension of 5 minutes at 72°C. We pooled the amplicons and size-selected the targeted fragments using a 2% Invitrogen™ UltraPure™ Low Melting Point Agarose gel (Fisher Scientific, Waltham, MA) followed by a purification using the Zymoclean Gel DNA Recovery Kit (Zymo Research, Irvine, CA). The libraries were sequenced at the Vanderbilt Technologies for Advanced Genomics using a NovaSeq 6000 System (Illumina, Inc, San Diego, CA) to produce pair-end reads 150 nucleotides long.

We processed the fastq files using an in-house bioinformatic pipeline similar to one published elsewhere (101). We only proceeded with reads #2 because it spanned our targeted DNA region. First, we used trimmomatic v.0.39 (102) to remove the sequencing adapters and filtered reads to retain those with a minimum length of 100 nucleotides and a minimum average quality score of 35. Then we used clumpify.sh from BBTools (https://sourceforge.net/projects/bbmap/) to remove duplicates. Lastly, we converted the file format from fastq to fasta using seqtk (103).

### High throughput long reads

We produced amplicons by assaying a PCR using the following oligonucleotides (forward: GGCAAGGAACTTGATGCACG and reverse: TGGCCAACCCACTGTTTGAT). The PCR reaction mix consisted of 0.2 IU/μl Phusion Hot Start II DNA Polymerase (Thermofisher Scientific, Waltham, MA), 1X Phusion HF Buffer, 200 μM dNTPs (Promega, Madison, WI), and forward and reverse oligonucleotides (IDT, Coralville, Iowa) at 0.10 μM each, in a final volume of 20 μl in 0.2 ml clear PCR tubes. The cycling conditions for this reaction were: 98°C for 1 minute, followed by 40 cycles of 98°C for 15 seconds, 55°C for 45 seconds, and 72°C for 1 minute, followed by a final extension of 4 minutes at 72°C. We confirmed the amplification by assaying 5 µl of each amplicon by electrophoresis on a 1.5% Agarose I™ gel before staining with Diamond™ Nucleic Acid Dye and imaging. When the amplification produced an amplicon, we used the remaining PCR products to prepare sequencing libraries with the Native Barcoding Kit 24 V14 (Oxford Nanopore Technologies, Lexington, MA) following the manufacturer’s instructions. We sequenced the libraries on a MinION Mk1C (Oxford Nanopore Technologies, Lexington, MA).

We processed the fast5 files with Guppy (v 6.4.2) (104) using the configuration file dna_r10.4.1_e8.2_260bps_sup.cfg for super high accuracy base calling. Next, we used porechop (https://github.com/rrwick/Porechop) to remove adapters and used Fitlong (https://github.com/rrwick/Filtlong) to remove sequences smaller than 500 nucleotides long and with a quality of less than 90%. We aligned the remaining sequences to the cattle reference genome (ARS-UCD1.2) using minimap2 (v 2.24) (105), allowing for spliced alignment (parameters: -ax map-ont --splice -c --cs=long --secondary=no --sam-hit-only -Y --splice-flank=no-G2k). Finally, we used samtools (106) to remove alignments with less than 500 nucleotides mapped to the genome and supplementary alignments.

### Sanger sequencing

We produced PCR amplicons using the procedures described for “High throughput long reads”. When the amplification produced an amplicon, we treated the remaining PCR products with 3 µl ExoSAP-IT™ Express PCR Product Cleanup Reagent (Thermofisher Scientific, Waltham, MA) and incubated at 37 °C for 15 minutes followed by 80 °C for 15 minutes. The sequencing assay was carried out by the Genomics Sequencing Center at Virginia Tech using the same forward oligonucleotide used for the initial PCR.

### Mapping and graphical visualization of DNA sequences relative to the bovine genome

For visualization and graphical representation of the edited sequences, we mapped resulting sequences in the fasta format to the cattle reference genome (ARS-UCD1.2) using the blat program (107) embedded in the UCSC genome browser (108, 109).

### PCR of a non-targeted region

For all samples failing to amplify with primers used targeted sequencing, we performed a PCR reaction targeting a segment of gene *CDK1* using the following oligonucleotides: GCCCAGACCCAGCATCATT, GGGAGTGCCCAAAGCTCTAAA (IDT). The reaction mix consisted of 0.2 IU/μl Phusion Hot Start II DNA Polymerase (Thermofisher Scientific, Waltham, MA), 1X Phusion HF Buffer, 200 μM dNTPs (Promega, Madison, WI), and forward and reverse oligonucleotides at 0.10 μM each, in a final volume of 20 μl in 0.2 ml clear PCR tubes. The cycling conditions for this reaction were: 98°C for 1 minute, followed by 40 cycles of 98°C for 15 seconds, 60°C for 30 seconds, and 72°C for 45 seconds, before a final extension of 4 minutes at 72°C. To check for the presence of DNA, 5 µl of each PCR product underwent electrophoresis on a 2% Agarose I™ gel before staining with Diamond™ Nucleic Acid Dye and imaging.

### DNA and RNA sequencing of single embryos

We collected embryos for DNA and RNA-sequencing on stage codes six or seven (110). We washed in PBS 0.1% BSA fraction V, followed by removal of the zona pellucida by exposure to EmbryoMax® Acidic Tyrode′s Solution and gentle pipetting. Once the zona pellucida was removed, we washed the embryos in PBS 0.1% BSA fraction V twice and collected them individually in microtubes in approximately 1 µl PBS 0.1% BSA fraction V. We placed the tubes on dry ice and stored the samples at −80°C.

We lysed the embryos by adding 10 µl of lysis solution, consisting of: 8.3 µl Luna Cell Ready Lysis Buffer 2X (New England Biolabs, Ipswich, MA), 0.66 µl Luna Cell Ready RNA Protection Reagent 25X (New England Biolabs, Ipswich, MA), 0.66 µl Luna Cell Ready Protease 25X, (New England Biolabs, Ipswich, MA), and 0.33µl RNasin® Plus Ribonuclease Inhibitor (40U/ µl) (Promega, Madison, WI). We incubated the solution on ice for 10 minutes, mixed by pipetting, then split the solution into two tubes. One tube, dedicated to DNA sequencing was further incubated at 37°C for 15 minutes, followed by the addition of one µl of Luna Cell Ready Stop Solution 10X (New England Biolabs, Ipswich, MA). We extracted DNA (111) by adding a solution of Sodium Acetate (5M) for a final concentration of 2.5 M, followed by the addition of 150 µl of Ethanol 100%. We stored the solution −20°C for over 15 hours and precipitated the DNA by centrifugation at 15,000xg for 20 minutes at 4°C. We washed the pellet twice with 150 µl Ethanol 70% and eluted it with nuclease-free water. The DNA was used as template for amplification of the targeted region using the oligonucleotides and procedures for “High throughput long reads”. Preparation of libraries, sequencing and processing of sequences were carried out as described above.

We extracted total RNA and from each ½ blastocyst using TRIzol reagent with Phasemaker Tubes for enhanced RNA purity and yield (112–115). RNA was stored in 70% ethanol at −80°C (112) until library preparation. We assessed RNA integrity of samples not used for sequencing with a 2100 Bioanalyzer Instrument (Agilent, Santa Clara, CA) and RNA 6000 Pico Kit (Agilent, Santa Clara, CA). These tests require the total volume of extracted RNA, therefore, only test samples were assessed to ensure the quality and rigor of our procedures. We amplified cDNA using a modified mcSCRB-seq protocol and produced libraries using the Illumina DNA Prep kit (Illumina, Inc, San Diego, CA) (112, 116). The libraries were sequenced at the Vanderbilt Technologies for Advanced Genomics using a NovaSeq 6000 System (Illumina, Inc, San Diego, CA) to produce approximately 30 million pair-end reads 150 nucleotides long per sample.

We aligned the RNA-sequencing data to the cattle reference genome (117) (ARS-UCD1.2/bosTau9) obtained from the Ensembl database (118, 119) using HISAT2 (v2.2.1) (120), followed by filtering with samtools (v1.17) (106, 121) to remove alignment less than 100 nucleotides long and with more than 5% mismatch nucleotides, plus removal of duplicates with biobambam2 (v2.0.95) (122). Next, we counted the sequences matching the reference annotation (Bos_taurus.ARS-UCD1.2.105.gtf) using featureCounts (v2.0.1) (123).

### Immunofluorescence of OCT4 and NANOG

We thinned each blastocyst’s zona pellucida by brief immersion in EmbryoMax® Acidic Tyrode′s Solution. Embryos were then washed and fixed in 4% paraformaldehyde solution for 15 minutes at room temperature. Next, we washed the embryos before transfer into permeabilization solution (0.25% Triton X-100) for 30 minutes, then blocking solution (10% horse serum, Thermofisher Scientific, Waltham, PA), for 1 hour at room temperature. We carried out concurrent staining for OCT4 and NANOG proteins by incubation of embryos and antibodies (mouse OCT4-alexafluor594 monoclonal antibodies, sc-5279 and mouse NANOG-alexafluor488 monoclonal antibodies, sc-374001, Santa Cruz Biotechnology, Dallas, TX) at 4°C for 24 hours and room temperature for 1 hour, respectively. Following washes, we placed the embryos in DAPI solution for 5 minutes. Embryos were individually placed into 5µl droplets of phosphate buffered solution (Thermofisher Scientific, Waltham, PA) submerged in mineral oil on a chambered coverglass (Thermofisher Scientific, Waltham, PA) for imaging.

### Fluorescence imaging

We evaluated fluorescence in our PZs or blastocysts using a Nikon Ti Eclipse fluorescence microscope (Nikon) coupled to X-Cite 120 epifluorescence illumination system and a DS-L3 digital camera using the cubes for DAPI (ex: 340-380nm, em: 435-485nm), GFP (ex:465-495nm, em:515-555nm) or mCherry (ex:532-587nm, em:608-683nm). The microscope was controlled by NIS-elements Imaging Software (v.5.02).

### Statistical analyses

The analytical procedures used to analyze differences in embryo development and differential transcript abundance, including the supplementary tables and pertinent graphs are available at: https://biase-lab.github.io/crispr_dart/

#### Assessment of differences in embryo development

We recorded the number of embryos that developed to the blastocyst stage and the number of putative zygotes with arrested development prior to blastocyst formation at 164-166 hpf and 188-190 hpf for each culture drop. Culture drop was considered biological replicate. We analyzed count data (success of blastocyst development or developmental arrest) using a general linear model with a binomial family, which results in logistic regression analysis, using the “glm” function from the R package “stats”. We used the number of blastocysts and the number of putative zygotes that failed to develop into blastocysts as the dependent variable, and the group (control, scramble, or Cas treated) was a fixed effect. The Wald statistical test was conducted with the function “Anova” from the R package “car” (124). Finally, we carried out a pairwise comparison using the odds ratio and two-proportion z-test employing the “emmeans” function of the R package “emmeans”. The null hypothesis assumed that the odds ratio of the proportion (*p*) of two groups was not different from 1 (*H*_0_: *p*_1_/*p*_2_ = 1). We adjusted the nominal P value for multiple hypothesis testing with the Bonferroni approach and inferred significance when adjusted P value< 0.05.

We analyzed data obtained from single embryo culture, with each embryo as a biological replicate, using the exact binomial test in R with the function “binom.test” (125). Significance was inferred if the P value < 0.05.

#### Assessment of differences in fluorescence

First, we calculated corrected total cell fluorescence (CTCF) using the standard formula: Integrated Density – (Area of selected cell X Mean fluorescence of background readings). We obtained the measurements necessary for the formula using the NIS-elements Imaging Software (v.5.02). Next, we fitted a linear model using the “lm” function of the R package “stats” where Log2(CTCF) was the dependent variable. Replicate and group (fluorescence protein mRNA or fluorescence protein mRNA + Cas13a and targeting sgRNA) were included as fixed effects. We assessed the significance of the variables using the “Anova” function of the R package “car”. Next, we tested the pair-wise significance of the two groups by a t-score test employing the “emmeans” function of the R package “emmeans”. The null hypothesis assumed that the difference between two averages (*x̅*) was not different from zero (*H*_0_: *x̅*_1_−*x̅*_2_ = 0), and significance was inferred at alpha = 0.05.

#### Differential transcript abundance

In R software (126, 127), we created one matrix with the read counts for all samples and retained genes classified as protein-coding and long non-coding DNA for downstream analysis. We calculated counts per million (CPM) using the function ‘cpm’ from the R package ‘edgeR’ (128) and retained genes if CPM>1 in 4 or more samples. We also calculated transcript per million as described elsewhere (129, 130) used for plotting the data. We estimated differential transcript abundance between edited and control blastocysts employing the quasi-likelihood negative binomial generalized log-linear model from the R package “edgeR” (128) and the Wald test from the R package “DESeq2” (131). We inferred statistically significant differences when False Discovery Rate (132) was less than 0.1 in both tests.

## Acknowledgments

We thank Select Sires for the donation of semen straws used in this research. This project was supported by Agriculture and Food Research Initiative Competitive Grant no. 2018-67015-31936 from the USDA National Institute of Food and Agriculture.

