## Appendix for "Ablation of *OCT4* function in cattle embryos by double electroporation of CRISPR-Cas for DNA and RNA targeting (CRISPR-DART)"

**This PDF file includes:**

Fig. 1S. Amplicons produced from embryos electroporated for the deletion of a segment of *OCT4*. 2

Fig. 2S. Example of alignment with nanopore long sequences (gray) showing the gapped alignment (blue). 3

Fig. 3S. Two examples of alignment with nanopore long sequences (gray) showing the gapped alignment (blue), including some minor variants of the deletions. 4

Fig. 4S. Patterns of RNA-sequencing data aligned to *OCT4*. 5,6

Movie. S1. Representative time lapse video of a non-electroporated control embryo. 7

Movie. S2. Representative time lapse video of a presumptive OCT4-/- embryo. 8

Table S1. Blastocyst developmental rates for the experiments comparing one versus two electroporation sessions with Cas9D10A. 9

Table S2. Contrasts between experimental groups for blastocyst yield at 164-166 hpf. 10

Table S3. Contrasts between experimental groups for blastocyst yield at 188-190 hpf. 11

Table S4. Blastocyst developmental rates for CRISPR-DART targeting *OCT4*. 12

Table S5. Contrasts between experimental groups for blastocyst yield in experiments with CRISPR-DART targeting OCT4. 13

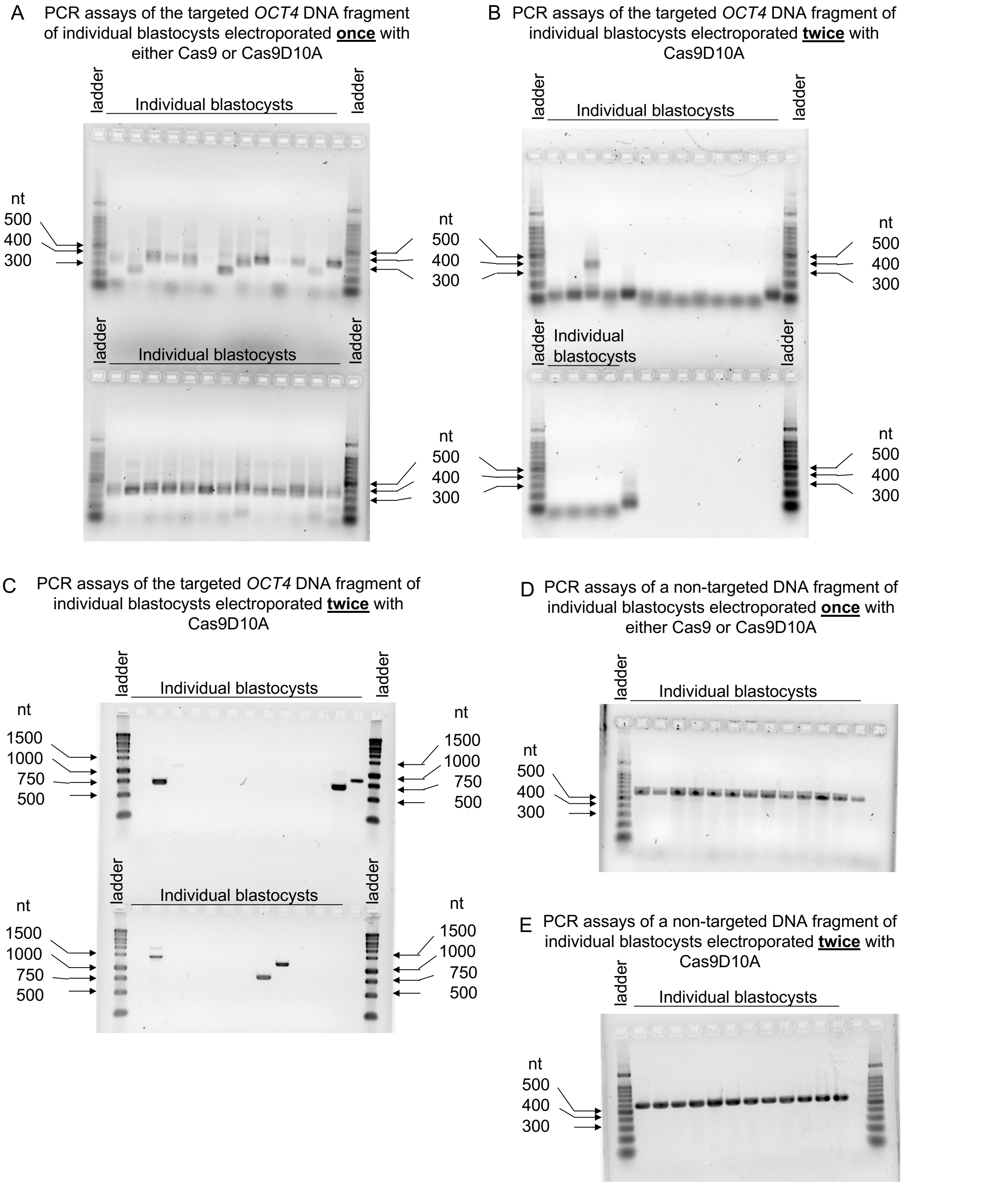


Fig. S1. Amplicons produced from embryos electroporated for the deletion of a segment of *OCT4*. (A) PCR assays of the targeted *OCT4* DNA fragment of individual blastocysts electroporated once with either Cas9 or Cas9D10A and sgRNAs. (B) PCR assays of the targeted *OCT4* DNA fragment of individual blastocysts electroporated twice with Cas9D10A and sgRNAs. (C) PCR assays of the targeted *OCT4* DNA fragment of individual blastocysts electroporated twice with Cas9D10A and sgRNAs. (D) PCR assays of a non-targeted DNA fragment of individual blastocysts electroporated once with either Cas9 or Cas9D10A and sgRNAs. (E) PCR assays of a non-targeted DNA fragment of individual blastocysts electroporated twice with Cas9D10A and sgRNAs.

Sample 8

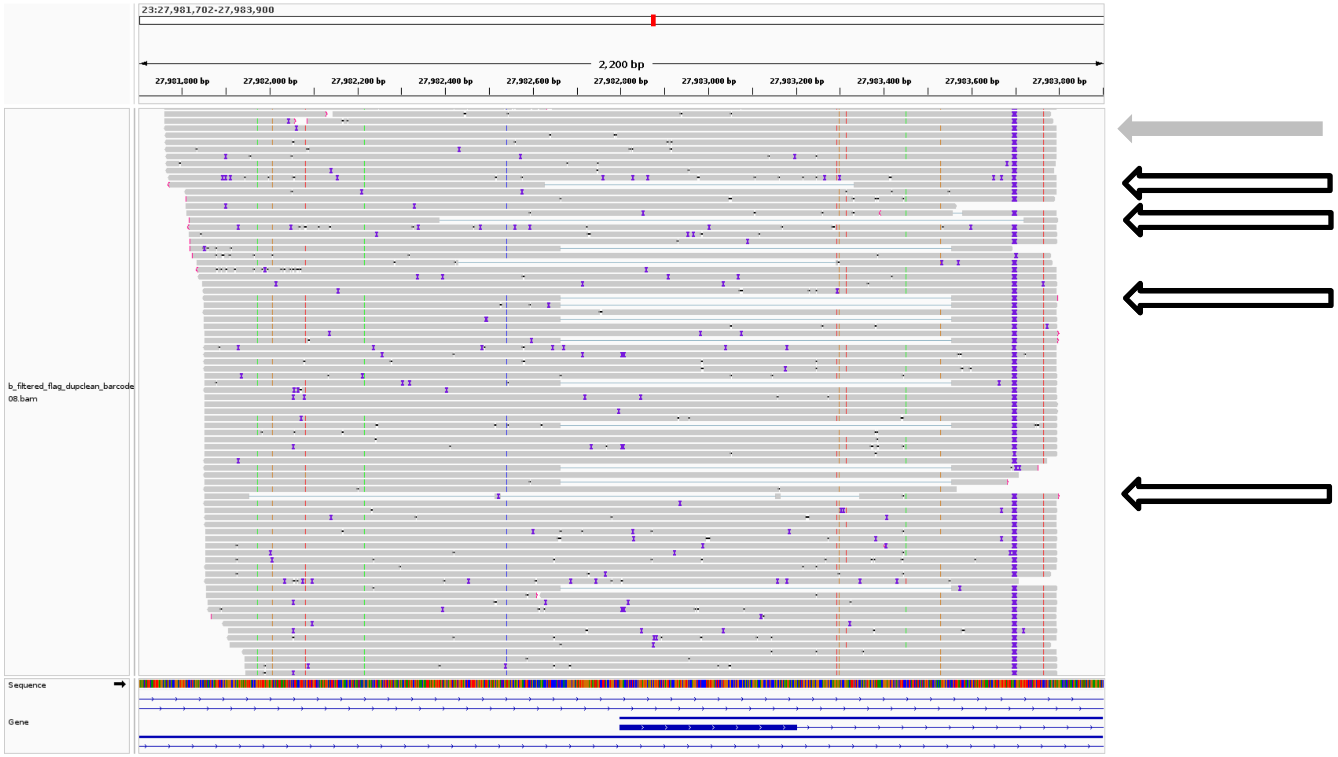


Fig. S2. Example of alignment with nanopore long sequences (gray) showing the gapped alignment (blue). This embryo was a mosaic with at least one wild-type chromosome sequence. Whole gray arrow point to one example of wild type sequence that covers the entire region of the chromosome targeted by the ribonucleoproteins. Empty arrows point to sequences that were produced from edited chromosomes by the ribonucleoproteins.

Sample 01

Sample 18

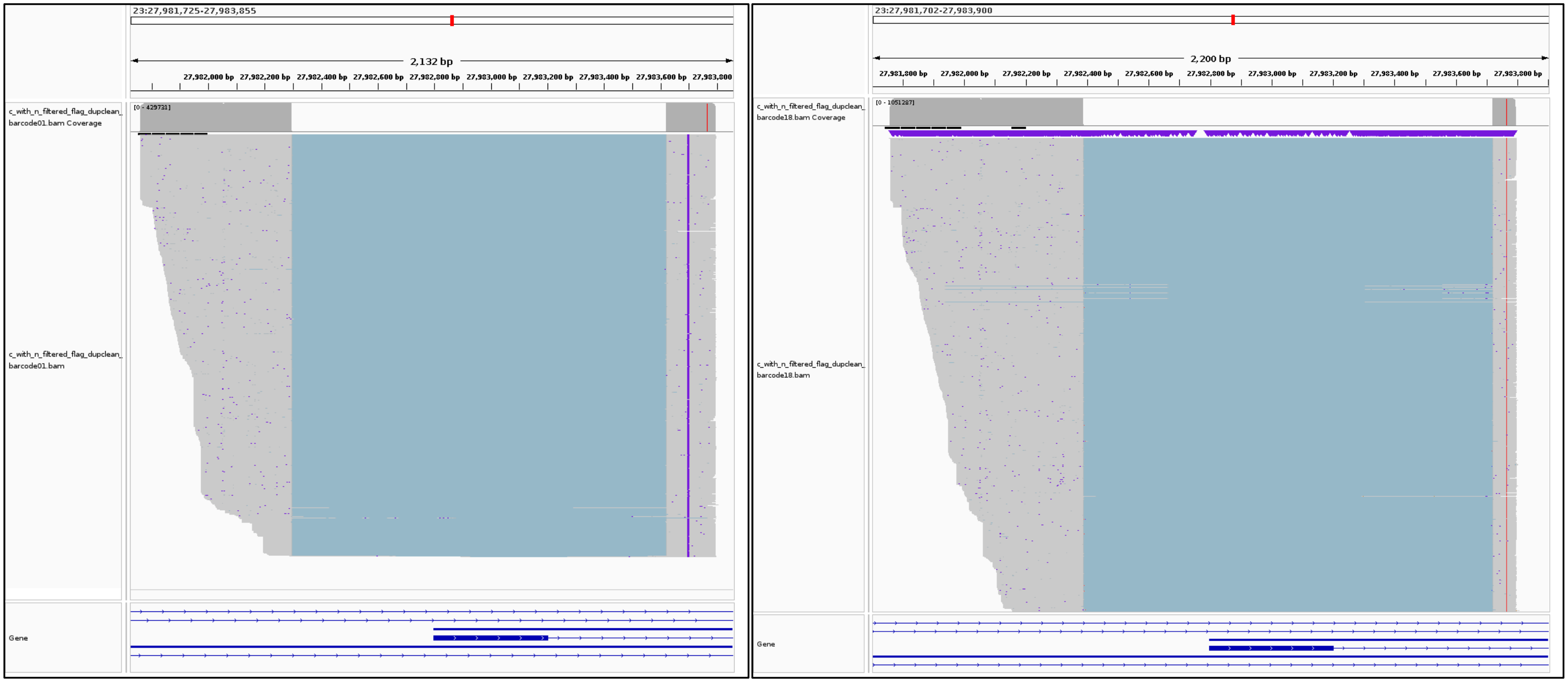


Fig. S3. Two examples of alignment with nanopore long sequences (gray) showing the gapped alignment (blue), including some minor variants of the deletions.

A

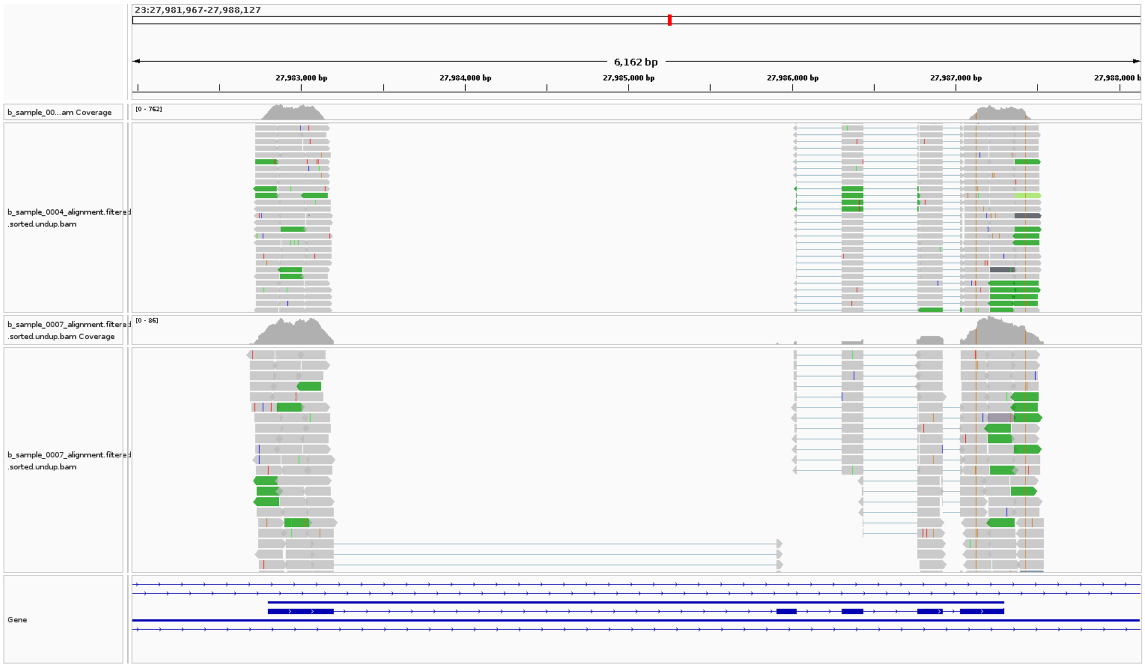


B

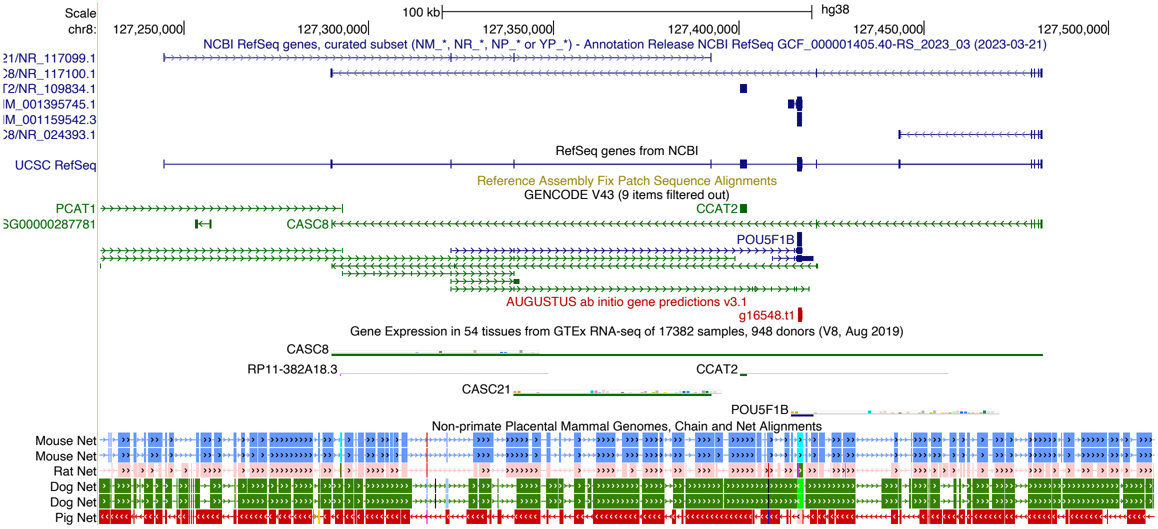


continue next page

C

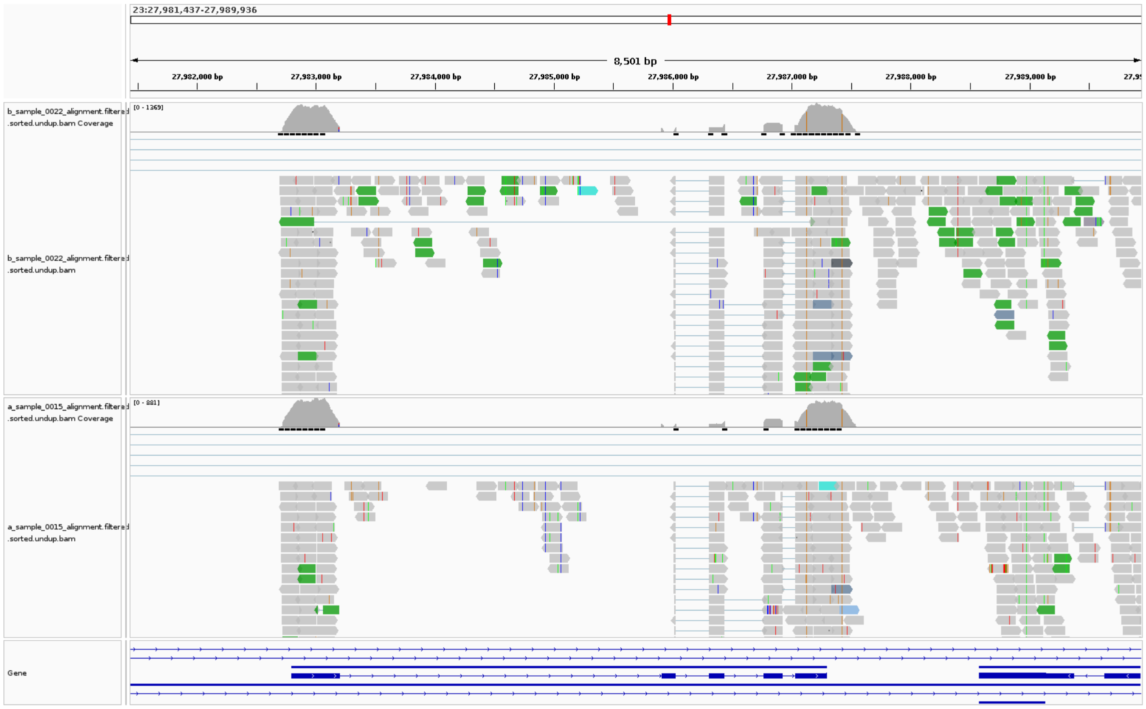


Fig. S4. Patterns of RNA-sequencing data aligned to OCT4. (A) Example of two blastocysts lacking exon 1 of the OCT4 gene. Note the absence of transcripts mapping to the introns. (B) Genome browser showing conservation of POU5F1b between human and other species. (C) Example of two wild type blastocysts. Note the presence of transcripts mapping to the introns.

Movie. S1. Representative time lapse video of a non-electroporated control embryo.

Movie. S2. Representative time lapse video of a presumptive OCT4-/- embryo.

Table S1. Blastocyst developmental rates for the experiments comparing one versus two electroporation sessions with Cas9D10A and sgRNAs.

| | | single electroporation | | double electroporation | |
|---|---|---|---|---|---|
| | | N PZ | % ± SE | N PZ | % ± SE |
| 164- | Cas9D10A + targeting sgRNAs | 337 | 6.8±1.4 | 655 | 3.2±0.7 |
| 166 | Cas9D10A + scramble sgRNAs | 146 | 17.1±3.1 | 209 | 17.7±2.6 |
| hpf | Controls not electroporated * | 182 | 25.3±3.2 | 182 | 25.3±3.2 |
| 188- | Cas9D10A + targeting sgRNAs | 337 | 11.6±1.7 | 655 | 7.9±1.1 |
| 190 | Cas9D10A + scramble sgRNAs | 146 | 31.5±3.8 | 209 | 28.2±3.1 |
| hpf | Control not electroporated * | 182 | 30.8±3.4 | 182 | 30.8±3.4 |

* The drops used for control culture (not electroporated) were carried out in parallel with the experiments of two versus one electroporation and thus used on statistical tests for single electroporation and double electroporation. hpf: hours post fertilization; N PZ: number of putative zygotes; SE: standard error

Table S2. Contrasts between experimental groups for blastocyst yield at 164-166 hpf.

| contrast | odds ratio | SE | DF | null | t ratio | raw P | BH P |
|---|---|---|---|---|---|---|---|
| Cas9D10A 1x / Scramble 1x | 0.3545 | 0.1092 | 15 | 1 | -3.3657 | 0.0042 | 0.0073 |
| Cas9D10A 1x / Control | 0.2166 | 0.0596 | 16 | 1 | -5.5584 | <.0001 | <.0001 |
| Scramble 1x / Control | 0.6109 | 0.1699 | 10 | 1 | -1.7722 | 0.1068 | 0.0849 |
| Cas9D10A 1x / Cas9D10A 2x | 2.2114 | 0.6847 | 31 | 1 | 2.5633 | 0.0154 | 0.0218 |
| Cas9D10A 2x / Scramble 2x | 0.1540 | 0.0441 | 26 | 1 | -6.5322 | <.0001 | <.0001 |
| Cas9D10A 2x / Control | 0.0979 | 0.0274 | 26 | 1 | -8.3042 | <.0001 | <.0001 |
| Scramble 2x / Control | 1.5723 | 0.3913 | 11 | 1 | 1.8185 | 0.0963 | 0.0838 |

1x: one electroporation session; 2x two electroporation sessions; SE: standard error; DF: degrees of freedom; BH P: Bonferroni corrected P values.

Table S3. Contrasts between experimental groups for blastocyst yield at 188-190 hpf.

| contrast | odds ratio | SE | DF | null | t ratio | raw P | BH P |
|---|---|---|---|---|---|---|---|
| Cas9D10A 1x / Scramble 1x | 0.2845 | 0.0701 | 15 | 1 | -5.1005 | 0.0001 | 0.0002 |
| Cas9D10A 1x / Control | 0.2945 | 0.0689 | 16 | 1 | -5.2232 | 0.0001 | 0.0002 |
| Scramble 1x / Control | 1.0350 | 0.2483 | 10 | 1 | 0.1434 | 0.8888 | 0.8888 |
| Cas9D10A 1x / Cas9D10A 2x | 1.5176 | 0.3390 | 31 | 1 | 1.8676 | 0.0713 | 0.0998 |
| Cas9D10A 2x / Scramble 2x | 0.2192 | 0.0463 | 26 | 1 | -7.1936 | <.0001 | <.0001 |
| Cas9D10A 2x / Control | 0.1940 | 0.0419 | 26 | 1 | -7.5892 | <.0001 | <.0001 |
| Scramble 2x / Control | 1.1299 | 0.2512 | 11 | 1 | 0.5496 | 0.5936 | 0.6925 |

1x: one electroporation session; 2x two electroporation sessions; SE: standard error; DF: degrees of freedom; BH P: Bonferroni corrected P values.

Table S4. Blastocyst developmental rates for CRISPR-DART targeting *OCT4*.

| | | N PZ | % ± SE |
|---|---|---|---|
| 164- | DART + targeting sgRNAs | 830 | 6.1±0.8 |
| 166 | DART + scramble gRNAs | 312 | 17.0±2.1 |
| hpf | Controls not electroporated | 348 | 23.0±2.3 |
| 188- | DART + targeting sgRNAs | 832 | 13.0±1.2 |
| 190 | DART + scramble gRNAs | 312 | 30.1±2.6 |
| hpf | Controls not electroporated | 348 | 31.6±2.5 |

hpf: hours post fertilization; N PZ: number of putative zygotes; SE: standard error

Table S5. Contrasts between experimental groups for blastocyst yield in experiments with CRISPR-DART targeting OCT4.

| | contrast | odds ratio | SE | DF | null | t ratio | raw P | BH P |
|---|---|---|---|---|---|---|---|---|
| 164- | DART + targeting sgRNAs / DART + scramble gRNAs | 0.3199 | 0.0668 | 35 | 1 | -5.4566 | <.0001 | <.0001 |
| 166 | DART + targeting sgRNAs / Control | 0.2193 | 0.0423 | 37 | 1 | -7.8746 | <.0001 | <.0001 |
| hpf | DART + scramble gRNAs / Control | 0.6855 | 0.1353 | 21 | 1 | -1.9129 | 0.0695 | 0.0834 |
| 188- | DART + targeting sgRNAs / DART + scramble gRNAs | 0.3460 | 0.0556 | 35 | 1 | -6.6000 | <.0001 | <.0001 |
| 190 | DART + targeting sgRNAs / Control | 0.3228 | 0.0499 | 37 | 1 | -7.3100 | <.0001 | <.0001 |
| hpf | DART + scramble gRNAs / Control | 0.9329 | 0.1575 | 21 | 1 | -0.4110 | 0.6852 | 0.6852 |

SE: standard error; DF: degrees of freedom; BH P: Bonferroni corrected P values.
